# The role of dynamic DNA methylation in liver transplant rejection in children

**DOI:** 10.1101/2022.05.03.489748

**Authors:** Mylarappa Ningappa, Xiaojian Shao, Chethan Ashokkumar, Qingyong Xu, Adriana Zeevi, Elin Grundberg, Tomi Pastinen, Rakesh Sindhi

**Author notes:** Co-Corresponding authors: Rakesh Sindhi, MD, FACS, Professor of Surgery, Co-Director, Pediatric Transplant Surgery, Director, Pediatric Transplant Research, University of Pittsburgh, Children’s Hospital of Pittsburgh of UPMC, Pittsburgh, PA 15201, USA, Tomi Pastinen, MD, PhD, Professor of Pediatrics, Director, Genomic Medicine Center, Children’s Mercy Research Institute, Children’s Mercy Kansas City. Authors contributed equally (Mylarappa Ningappa, Xiaojian Shao).

## Abstract

**Background:** Transcriptional regulation of liver transplant (LT) rejection may reveal novel predictive and therapeutic targets.

**Purpose:** To test the role of differential DNA methylation in children with biopsy-proven acute cellular rejection (rejectors, R) after LT.

**Methods:** Paired peripheral blood DNA samples were obtained before and after LT from 17 children, including 4R and 13 non-rejector (NR), and assayed with MethylC capture sequencing (MCC-Seq) approach covering 5 million CpGs in immune-cell specific regulatory elements. Differentially methylated CpGs (DMCs) were identified using generalized linear regression models adjusting for sex and age and merged into differentially methylated regions (DMR) comprising 3 or more DMCs.

**Results:** Contrasting R vs NR, we identified 2238 DMCs in post-LT and 2620 DMCs in pre-LT samples, which clustered in 216 and 282 DMRs respectively. DMCs associated with R were enriched in enhancers and depleted in promoters. The proportion of hypomethylated versus hypermethylated DMRs increased from 22% to 48% (p<0.0001) in pre-LT vs. post-LT DMCs, respectively. The highest-ranked biological processes enriched in post-LT DMCs were antigen processing and presentation via MHC class I, MHC class I complex, and peptide binding (p<7.92E-17), respectively. Top-ranked DMRs mapped to genes which mediate B-cell receptor signaling (*ADAP1*) or regulate several immune cells *(ARRB2)* (p<3.75E-08). DMRs in MHC class I genes were enriched for SNPs which bind TFs, affect gene expression and splicing, or alter peptide-binding amino acid sequences.

**Conclusions:** Dynamic methylation in distal regulatory regions reveals known transplant-relevant MHC-dependent rejection pathways, and identifies novel loci for future mechanistic evaluations in pediatric transplant subcohorts.

## Introduction

Epigenetic changes may be better suited to aid management of liver transplant (LT) rejection in children, who are at risk for cumulative toxicity of lifelong immunosuppression.^1^ These changes regulate gene transcription by affecting the binding of transcription factors (TF) and should therefore precede transcription and its clinical consequences. In turn, patterns of TF binding or the loci which bind to these factors can reveal the particular mechanism of rejection in a given recipient, and the immunosuppressant best able to target that mechanism. This task is not fulfilled by currently available diagnostic tests. Cell function assays predict rejection-risk and not the appropriate immunosuppressant.^2^ Molecular diagnostics detect rejection and its progression using predominantly upregulated genes, which may not be mechanistic.^3,4^

As a clinically usable epigenetic mechanism, DNA methylation is attractive for several reasons. Methylation status is correlated with chromatin states, which require complex sample preparation procedures for their characterization.^5^ As a test substrate, DNA can be easily acquired from several sources: blood, biopsy or body fluids. Commercially available arrays use small quantities of DNA to screen methylation status of nearly a million methylation (CpG) sites genome-wide.^6^ Early DNA methylation studies have identified differentially methylated regions near genes encoding cytokines as well as other genes within interferon and mTOR signaling pathways, as being relevant in renal transplant rejection. ^7,8^ No such work has been performed in pediatric LT recipients.

We and others have shown that trait-associated and dynamic epigenetic variants are enriched in distal regulatory enhancer regions and we further showed that these features are also seen downstream of the transcription start sites (TSS)– both genomic regions being underrepresented on commercially available arrays.^5-11^ To overcome this limitation, we implemented the methylC-capture sequencing (MCC-Seq) approach for customized DNA methylation profiling of millions of CpGs located in regulatory elements known to be active specifically in immune cells.^11,12^ Specifically, this immune cell MCC-Seq panel covers (1) the majority of human gene promoters, blood-cell-lineage-specific enhancer regions and methylation footprint regions observed in peripheral blood,^12^ (2) CpGs from Illumina Human Methylation 450 Bead Chips, and (3) published autoimmune-related SNPs as well as SNPs in their linkage disequilibrium regions with r^2^ > 0.8. Here we test whether MCC-Seq applied to pre- and post LT blood samples obtained in the first 90 days after LT from 17 children with LT can reveal potential mechanisms of LT rejection for further investigation. Four children experienced early biopsy-proven acute cellular rejection (R) and 13 did not (NR).

## Methods

### Human subjects

Archived DNA extracted from blood samples from 17 children with LT was tested under University of Pittsburgh Institutional Review Board approved study #19030279. Mean±SD age of subjects was 6.3±8.2 years and male: female gender distribution was 6:11. Four of the 17 children experienced acute cellular rejection within 90 days of transplant and were termed rejectors (R). Thirteen of the 17 children had no rejection (NR). For the four rejectors in the post-LT cohort, samples were collected at 9.75±10.8 days before the rejection event.

### MethylC-capture sequencing (MCC-Seq)

One µg of DNA was used for whole genome bisulfite sequencing library preparation (KAPA Biosystems, Wilmington, MA), bisulphite conversion (Epitect Fast DNA Bisulfite Kit (Qiagen)), and enrichment (12-plex) using the custom probes ^10,13^ according to the Roche NimbleGen SeqCapEpi Enrichment System protocol (Wilmington, MA). We compared DNA methylation differences between R and NR using a general linear regression model (GLM) adjusted for age and sex. Correction of immune cell proportion was performed with constrained linear projection via the projectMix function of the RefFreeEWAS package (version 2.2), using a custom panel of 30,455 cell-type specific hypomethylated and hypermethylated CpGs.^14^ The blood reference epigenome profiles include Megakaryocyte, Neutrophil, Monocyte, B-cell and T-cell. Differentially methylated CpGs (DMCs) were further filtered if the DNA methylation profile is correlated with any of the estimated blood proportion at nominal p-value < 0.05. To assess potential regional clustering of significant DMCs, candidate regions surrounding the blood corrected DMCs were expanded for up to 200bp distance both upstream and downstream. Within these candidate regions, all consecutive CpGs with methylation changes in the same direction and with nominal p-value <0.01 were merged. Regions with at least 3 CpGs fulfilling these criteria were considered differentially methylated regions (DMRs). GO function enrichment analysis of rejection associated DMCs was performed using Genomic Regions Enrichment of Annotations Tool (GREAT).^15^

## Results

### CpGs that characterize rejectors are enriched in distal regions and hypomethylated after LT

Of the ∼2,5 million autosomal CpGs tested in the association analysis, we identified 3,357 DMCs including 1,894 hypo-methylated and 1463 hyper-methylated CpGs in post-LT samples from R compared with NR. The genome-wide distribution of significant CpGs is shown in Figure 1A. Among the 3357 DMCs, 1117 were significantly correlated with blood cell proportions and were excluded. Among the remaining 2,238 DMCs, 1,108 were hypermethylated and 1,130 were hypomethylated (Tables 1 and Table S1). To further explore how the DMCs mapped to regions, we identified differentially methylated regions (DMRs) containing clusters of 3 or more DMCs with the same directional change in methylation. Of 216 DMRs that clearly distinguished R and NR groups, 113 were hypermethylated and 103 were hypomethylated (Figure 1B, Table S2). Examples include intronic regions in *ADAP1* (50 DMCs, chr7:948678-949115, Figure 1C) and *LHX6* (34 DMCs), and the intergenic region near *ARRB2* (24 DMCs) (Table S2). ^16-19^ Enrichment analysis of genomic regions revealed that rejection associated DMCs and DMRs were enriched in the distal regulatory regions and depleted in the promoter regions (Table S2). Next, we identified significant genome-wide DMCs in pre-LT samples from R compared with NR subjects. After filtering DMCs (n=665) which correlated significantly with blood proportions, 2,620 DMCs remained of which 1413 were hypermethylated and 1207 were hypomethylated (Table S3). Among these DMCs, we identified 282 DMRs consisting of 221 hypermethylated and 61 hypomethylated DMRs (Table S4).

**Table 1.**
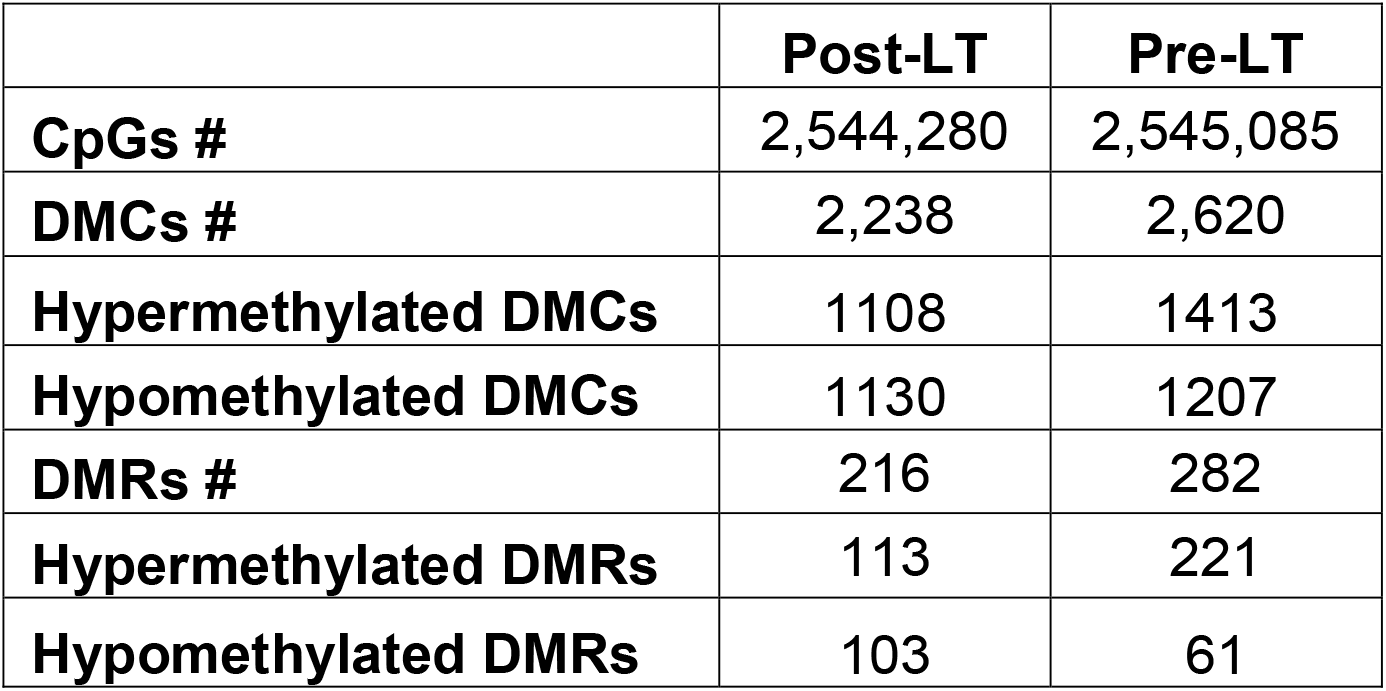
Counts of differentially methylated CpGs (DMCs) (Table S1, Table S3), and regions (DMRs) (Table S2, Table S4), in samples obtained from 4 rejectors (R) compared with those from 13 non-rejectors, within 90 days after (post-LT) and before (Pre-LT).

|  | **Post-LT** | **Pre-LT** |
| --- | --- | --- |
| **CpGs #** | 2,544,280 | 2,545,085 |
| **DMCs #** | 2,238 | 2,620 |
| **Hypermethylated DMCs** | 1108 | 1413 |
| **Hypomethylated DMCs** | 1130 | 1207 |
| **DMRs #** | 216 | 282 |
| **Hypermethylated DMRs** | 113 | 221 |
| **Hypomethylated DMRs** | 103 | 61 |

**Figure 1.**
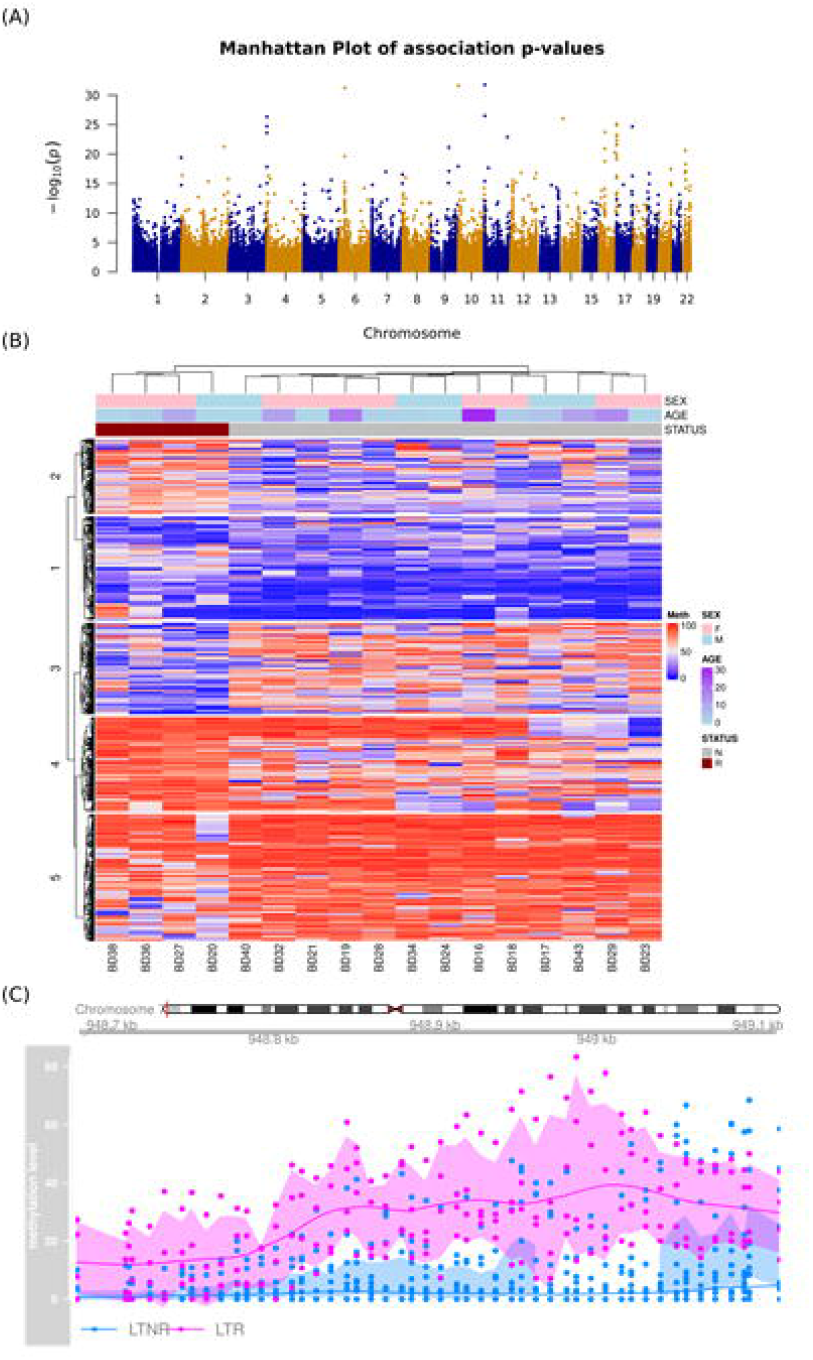
Identification of differentially methylated CpGs (DMCs) for LT rejection. **(A)** Manhattan plot of p-values from association analysis to identify differentially methylated CpGs **(B)** Heatmap of DMCs in methylation profiles for all individuals. Phenotype features including sex, age onset, and LT rejection status) are illustrated in the bars on top of the heatmap. **(C)**. DNA methylation pattern at the (differentially methylated region (DMR) for *ADAP1* (chr7:948678-949115). This region shows consistent hyper-methylation among R (pink) compared with NR (blue) over 50 DMCs.

### Functional enrichment analysis reveals mechanisms for additional investigation

The genes associated with DMCs located within the DMRs regions in post-transplant samples were enriched for the biological processes of “antigen processing and presentation of exogenous peptide antigen via MHC class I, (p-value=4.4e-21) and cellular component of “MHC protein complex” (p-value=1.7e-23) and molecular function of peptide antigen binding (P value =7.9E-17) (Table S5). DMCs in pre-transplant samples were enriched for the top-ranked biological processes of “regulation of thyroid-stimulating hormone secretion” (p-value<3.1e-58) and several developmental and morphogenesis processes. “Antigen processing and presentation of exogenous peptide antigen via MHC class’’ ranked 17^th^ among these processes (p-value <9.5e-34) (Table S6). Enriched top-ranked cellular components included “MHC class I protein complex” (p-value <9.1e-36), and top-ranked molecular functions included “thyroid-stimulating hormone receptor activity” (p-value<2.6e-55)”, and “peptide antigen binding” (p-value <1.8e-30) (Table S6). Enrichment of DMRs in MHC class I molecules among rejectors was demonstrable in coding sequences of *HLA-C* and intronic regions of *HLA-F* in pre-transplant samples and the *HLA-B* promoter sequence in post-transplant samples.

Public databases reveal that all three HLA DMRs (*HLA-C, HLA-F* and *HLA-B*) contain known single nucleotide polymorphisms, which are known to bind to transcription factors, alter expression of the corresponding gene, or affect peptide antigen binding (SNPs) (Table S7 A-C, Figure S1). SNPs associated with altered gene expression or expressed quantitative trait loci are shown in Figure S1. However, the differential methylation between R vs NR was not associated with the degree of mismatches between the donor and recipient in traditional pre-transplant HLA typing of HLA-B (mean±SD 1.5±0.58 vs 1.69±0.48, p=0.577, (Not significant, NS) and HLA-C alleles (mean±SD 1±0.82 vs 1.38±0.51, p=0.42, NS).

## Discussion

Our study shows that rejection after LT in children is associated with differential DNA methylation affecting distal regions of genes, and with 3-fold increase in hypomethylated: hypermethylated regions (103:113) compared with pre-transplant status (61:221, p=0.0001) (Figure 1B, Table 1). This is to be expected because dynamic changes in methylation which are induced by disease predominantly affect distal regions such as enhancers, for which the MCC-seq platform is especially suited. Differentially methylated loci also reveal several new directions for future study. *First*, DMRs containing the highest number of DMCs are present in several loci which could regulate the rejection alloresponse (Table S2). These loci include *ADAP1* (Figure 1C) which participates in B-cell receptor signaling;^17^ *ARRB2, which* negatively regulates inflammatory responses of many immune cells including T- and B-cells;^18^ and *LHX6* which regulates the development of many cells including lymphoid cells.^19^ A larger association study and functional experiments are needed to confirm this possibility.

Second, enrichment analyses suggest that post-transplant rejection may recruit MHC I molecules, which classically present endogenous antigen, to present exogenous (transplant) antigens.^20^ Exogenous antigens are usually presented by MHC class II molecules. This possibility is based on the emergence of TAP-independent antigen processing and presentation of peptide antigen via MHC class I, as the top-ranked biological process, and MHC class I protein complex and peptide antigen binding as the top-ranked cellular component and molecular function, respectively, in post-transplant samples from rejectors compared with non-rejectors (Table S5). The transporter associated with antigen processing (TAP) proteins bind to the complex between MHC class I molecules and endogenous peptide antigens, aiding transport of this complex to the cell surface.^16,20^ Presentation of exogenous antigen via MHC class I does not require TAP. Consistent with a shift to TAP-independent antigen presentation after transplantation, pre-transplant DMCs are enriched for molecular functions of TAP1-TAP2-binding and peptide antigen binding, and the cellular component, MHC class I protein complex. Pre-LT samples were also enriched for several biological processes related to morphogenesis, for example pronephric field specification and nephron morphogenesis, reflecting the contribution of underlying developmental diseases that require liver transplantation, to the pre-transplant blood methylome (Table S6).

Third, these enrichment results suggests that dynamic methylation could add to the clinical utility of the MHC locus by predicting transplant rejection. Currently, donor-recipient mismatches at polymorphic HLA alleles are used to assess histocompatibility. These mismatches lead to donor-specific anti-HLA antibodies (DSA), which are increasingly implicated in late graft loss, and DSA specificity. ^20-25^ Highly polymorphic HLA genes, for which corresponding proteins show high expression on the cell surface induce DSA more frequently (*HLA-DR, HLA-DQ*, and *HLA-B*). ^26,27^ Polymorphisms in *HLA-B*, which is enriched for DMRs in post-transplant samples from rejectors are also associated with TAP-independent MHC class I antigen processing and presentation of exogenous antigens and have been used to explain CD8-mediated cytotoxicity in a variety of autoimmune diseases.^28-31^ TAP-independent class I antigen presentation after LT may also explain why alloresponsive CD8-memory cells are also being used clinically to predict acute liver transplant rejection in children.^2^ Others HLA genes are less polymorphic and have low surface expression because of predominantly intracellular location, which is influenced in part by binding to TAP. These genes are either associated with a low incidence of DSA (*HLA-C*) or have no known function *(HLA-F). HLA-C* is largely intracellular where it remains bound to TAP.^26^ *HLA-F* expression is restricted to the B-cell lineage in the resting state, also partly influenced by binding to TAP.^27^ DMRs in *HLA-C* and *HLA-F* are enriched in pre-transplant samples from rejectors in our study.

Finally, public databases reveal how sequence variants in DMR of the HLA genes may affect the rejection alloresponse via dynamic methylation, highlighting a complexity that has yet to be understood fully. The expression of MHC molecules is regulated by binding of TFs to regulatory elements.^32-34^ This binding is influenced by polymorphisms at TF binding sites. An example is the CCCTC binding transcription factor, CTCF, which regulates the expression of several MHC genes.^35,36^ We have previously shown the effect of altered methylation on CTCF-mediated expression of *HLA-DOA*, a B-cell-specific MHC gene, and the relationship of SNPs in these regulatory loci to B-cell presentation of donor antigen during LT rejection in children.^37,38^ Consistent with these reports, each DMR in the *HLA-B, -C* and *-F* genes contains known SNPs, which either bind TFs, bring about amino acid changes in the peptide-biding regions of corresponding MHC molecules, or may explain cell-cell interactions such as those between HLA-F and natural killer cells, which underlie immunity and tolerance (Table S7). For example, the SNPs rs2076177 and rs1736924 in the DMR of *HLA-F* in rejectors are respectively associated with enhanced and decreased expression and splicing in public databases (Figure S1). Thus, the study of dynamic DNA methylation can reveal potential causal links between genetic variation and transcriptional regulation and facilitate preventive intervention by predicting rejection.

In conclusion, and consistent with several previous studies in non-transplant settings, our study localizes dynamic DNA methylation associated with LT rejection in children, to mostly non-promoter distal regulatory regions. The emergence of the well-known transplant-relevant MHC-dependent antigen presentation as an enriched pathway, and of associated MHC class I-dependent mechanisms among rejectors in this exploratory cohort adds to the validity of DNA methylation as an investigative tool. Given that this enrichment is associated with novel but plausible differentially methylated regulatory loci, DNA methylation analysis can identify novel mechanisms and hypotheses for further investigation in the rare pediatric liver transplant sub-cohort of transplant recipients.

## Abbreviations

ADAP1: ArfGAP With Dual PH Domains 1
ARRB2: arrestin B2
CTCF: CCCTC binding transcription factor
DMC: differentially methylated CpG
DMR: differentially methylated regions
GLM: generalized linear regression models
LHX6: LIM Homeobox 6
LT: Liver transplantation
MCC-Seq: methylation capture sequencing
MHC: major histocompatibility complex
mTOR: mammalian target of rapamycin
NR: non-rejectors
R: Rejectors
SNP: single nucleotide polymorphism
TF: transcription factor
TSS: transcription start site

---

**Figure S1:** SNPs in DMRs that are known to affect HLA gene expression (eQTL) or splicing (sQTL) of the corresponding gene. Figures adapted from https://gtexportal.org/. SNP=single nucleotide polymorphism, DMR=differentially methylated region.

