## Supplementary figures and images for "The role of dynamic DNA methylation in liver transplant rejection in children"

### Figure S1

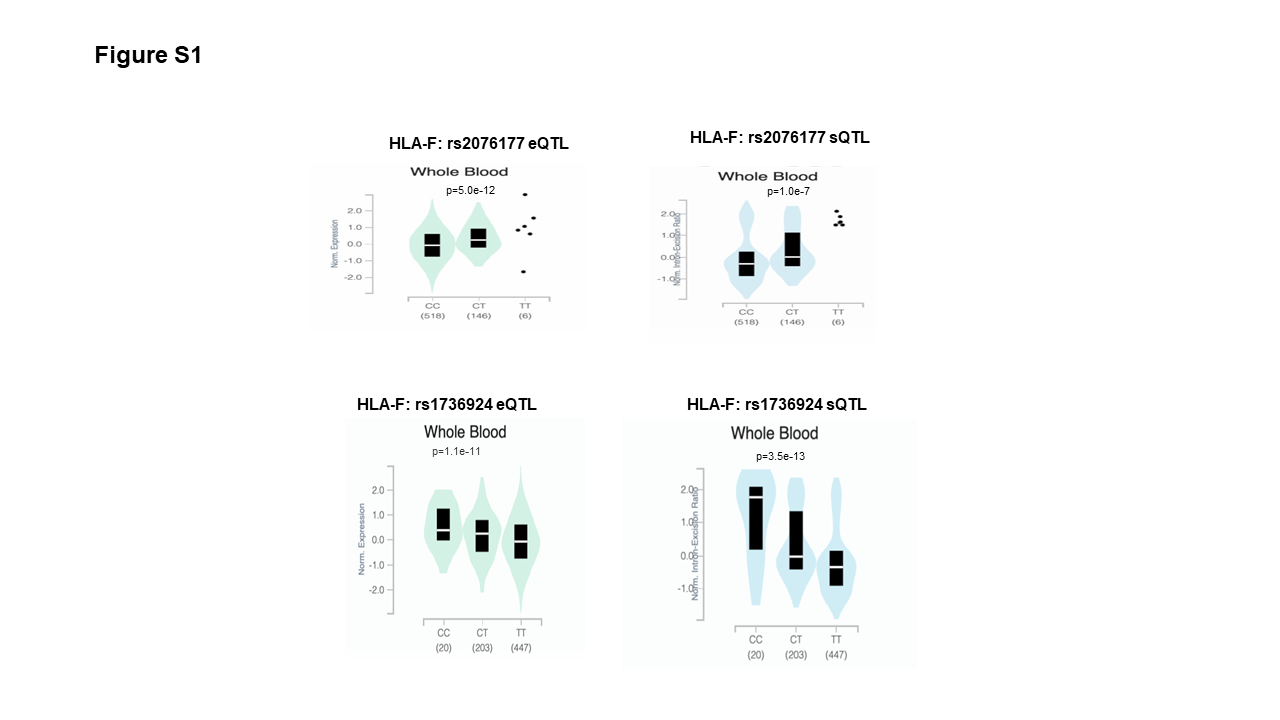
